# Cystic fibrosis transmembrane conductance regulator (CFTR) downregulates the excitability of deep pyramidal neurons in the rodent cortex

**DOI:** 10.1101/2023.08.29.555433

**Authors:** Zongyue Cheng, Xuan Sun, Wang Xing, Fei Luo, Hsiao Chang Chan, Wenbiao Gan, Baoming Li

## Abstract

The cystic fibrosis transmembrane conductance regulator (CFTR) is an ATP-binding chloride channel that is regulated by intracellular cAMP/PKA phosphorylation. CFTR is widely expressed in peripheral tissues and organs of mammals and plays a vital role in maintaining chloride balance and cellular homeostasis. While preliminary studies have identified CFTR expression in the central nervous system (CNS), it is not clear whether this channel also modulates the neural network of the cerebral cortex by regulating the intracellular chloride level of neurons. In this study, we employed immunohistochemical staining, patch-clamp recording, and two-photon imaging techniques to comprehensively analyze the functions of the CFTR channel in the rodent cortex. Our results indicate that CFTR is primarily distributed in the deep pyramidal somata and superficial axons of the cerebral cortex. Regulation of CFTR has the potential to alter the resting membrane potential and evoke action potentials of layer V pyramidal neurons, which produces significant changes in inhibitory synaptic transmission. Furthermore, we found that inhibiting CFTR channels increased the calcium activity of axon boutons and somata of the primary motor cortex in vivo, promoting motor learning. Overall, these findings implicate a crucial role of CFTR in modulating Cl-homeostasis and neuronal excitability in the cerebral cortex, furthering our understanding of the functions of the chloride channel in the central nervous system.

## 1. Introduction

Cystic fibrosis (CF) is the most common autosomal recessive genetic disease among Northern European ancestry^1–5^. It usually manifests as persistent pneumonia, intestinal obstruction, pancreatic insufficiency, and male infertility^6–13^. The mutation of ATP-binding superfamily member cystic fibrosis transmembrane conductance regulator (CFTR) results in CF^1,5^. CFTR is widely expressed in the epithelial tissues of the respiratory, digestive, and reproductive systems^14–21^. The CFTR channel can permeate Cl^-^ and HCO_3_ ^-^; coordinate the transfer of water and HCO_3_ ^-^ of epithelial tissues and regulate cell pH and water-salt balance^22^.

Recent studies have shown that CFTR is widely expressed in both the peripheral and central nervous systems^23–25^. Specifically, CFTR has been found to be concentrated in various sympathetic ganglia and dorsal root ganglia ^26–28^, while also being more uniformly distributed throughout regions of the brain such as the frontal lobe, parietal lobe, temporal lobe, occipital lobe, thalamus, hippocampus, pons, cerebellum, and medulla oblongata^29–32^. Additionally, the presence of CFTR mRNA has been detected in the spinal cord^33–35^. Despite the lack of a uniform phenotype among cystic fibrosis patients, it has been observed that they are disproportionately affected by central nervous system issues compared to the general population^36–40^. This suggests a possible link between CFTR expression and nervous system pathology in cystic fibrosis.

While the function of CFTR in epithelial cells has been thoroughly investigated, its role in the nervous system remains largely unknown. In newborn rats, CFTR has been found to contribute to chloride homeostasis and neuronal excitability in spinal motoneurons and interneurons^34,41^. Additionally, CFTR has been shown to exert a de-inhibitory regulatory effect on postsynaptic glycine inhibition in the trigeminal motor nucleus during rapid eye movement sleep^33^. These findings suggest that CFTR may function as an ion channel in the nervous system to maintain cell homeostasis and chloride balance^42,43^. However, there is currently a lack of evidence to support the involvement of CFTR in chloride channel regulation and neuronal excitability in the cerebral cortex. In this study, we employed various techniques such as immunohistochemistry, patch-clamp recording, and two-photon imaging to systematically investigate the function of CFTR in the rodent cerebral cortex. Our findings provide insight into the role of CFTR in modulating the excitability of pyramidal neurons in the deep layer of the cerebral cortex, as well as its impact on the learning process of live animals. Our research has important theoretical implications for understanding the mechanisms and functions of CFTR in the cerebral cortex.

## 2. Results

### CFTR is expressed on the somata and axons of pyramidal neurons in the deep layer of the cerebral cortex

Previous research has demonstrated that the expression levels of CFTR are similar across various regions of the brain. To determine the specific cell type in which CFTR is present in the cerebral cortex, we utilized two validated CFTR antibodies (ab117447 and ACL-006, Supplementary Figure 1) to stain the cerebral cortex of rats. Our findings indicate that CFTR is highly expressed in the somata and processes of neurons (Supplementary Figure 2). Through immuno-double labeling, we discovered that CFTR is primarily found in layer V/VI of the cerebral cortex, in conjunction with CaMK II, a marker for excitatory neurons (Fig 1a-d, Supplementary Figure 3). CFTR also shows significant co-localization with the mature axon marker NF200 (Fig. 1e-i). This distribution pattern was consistent in cultured mature neurons as well (Supplementary Figure 4).

**Figure 1.**
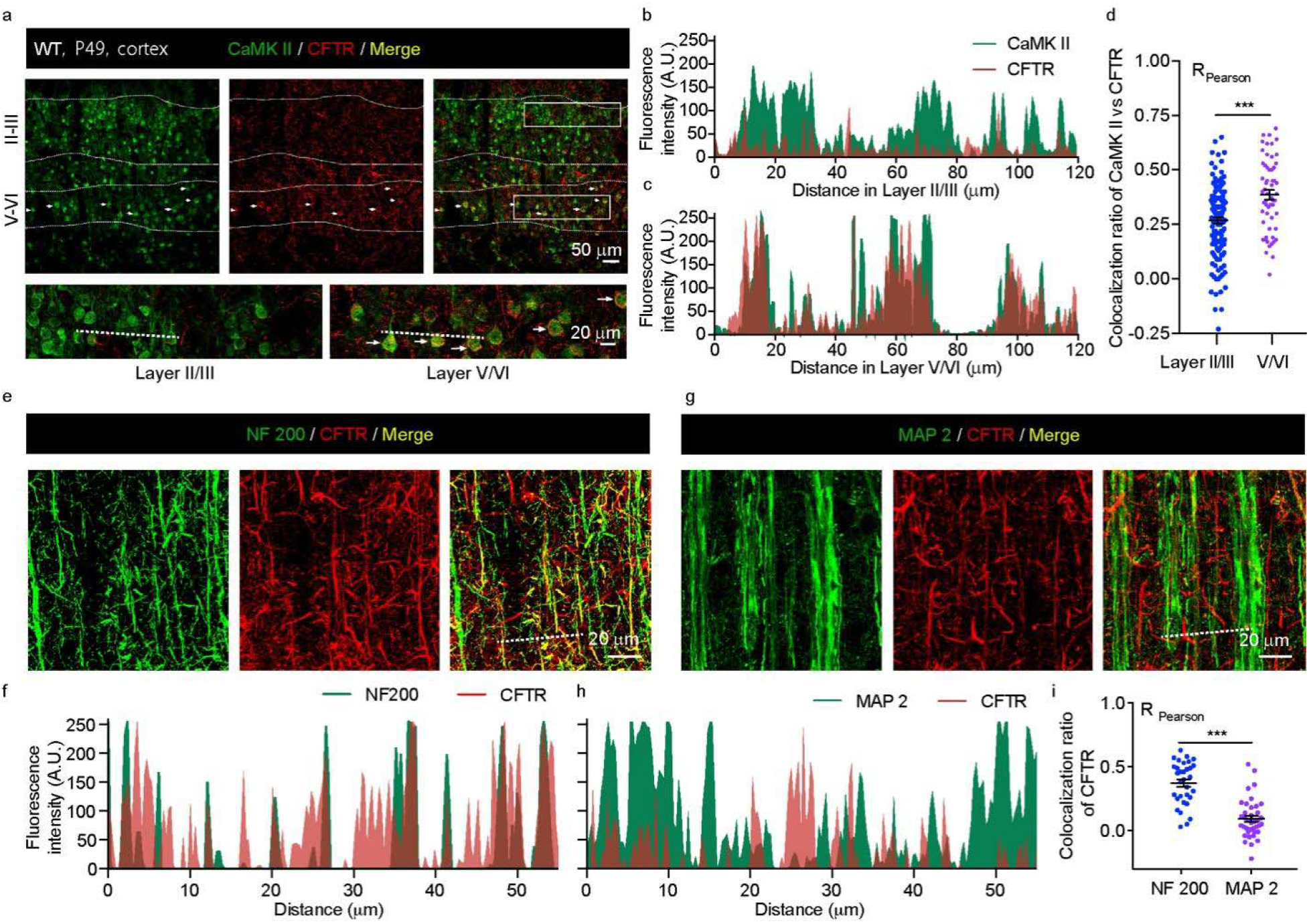
CFTR is expressed in the somata and axons of pyramidal neurons in the deep layer of the cerebral cortex. Confocal laser microscopy shows the staining results of cerebral cortex sections of adult rats (P49). The nucleus (DAPI) is colored blue, CFTR is colored red, and other cell markers are shown green. **a**. Double immunolabeling results of CFTR and excitatory neuron marker CaMK II. The white arrows indicate somata with a good co-standard. **b**. The fluorescence distribution of CaMK II and CFTR in the superficial cortex (Layer II/III) along the dotted line in Figure a. **c**. The fluorescence distribution of CaMK II and CFTR in the deep cortex (Layer V/VI) along the dotted line in Figure a. **d**. Pearson co-localization coefficient of CFTR and CaMK II in the superficial and deep layer. -1 to 1 indicate the interval of the co-localization relationship. It is generally believed that there is a moderate or above co-localization relationship when R (Pearson) is greater than 0.4 indicates, and when R (Pearson) is less than 0.2, it is assumed that there is no clear co-localization relationship. **e**. Double immunolabeling results of CFTR and mature axon fibers (NF200) in layer I of the cerebral cortex. **f**. the fluorescence distribution of NF 200 and CFTR along the dotted line in Figure e. **g**. Double immunolabeling results of CFTR and dendritic fibers (MAP2) in the superficial cerebral cortex. **h**. the fluorescence distribution of MAP2 and CFTR along the dotted line in Figure g. **i**. Pearson commonness coefficient of NF200 and MAP2. **d, i**, Data are mean ±s.e.m. *** *P* < 0.001. n =

To solidify the existence of CFTR in the deep layer of the cerebral cortex, we utilized a whole-cell patch-clamp technique to observe CFTR activity stimulated by a step voltage in the frontal cortex of rats (Fig. 2a). By using Gly H-101, a specific inhibitor of CFTR, we were able to decrease Cl-currents that were mediated by the cAMP complex in pyramidal neurons (Fig. 2b-d). Additionally, the CFTR activator RP-107 increased the chloride current in cell-attached cells, which was suppressed by the CFTR inhibitors Glibenclamide and Gly H-101 (Supplementary Figure 5).

**Figure 2.**
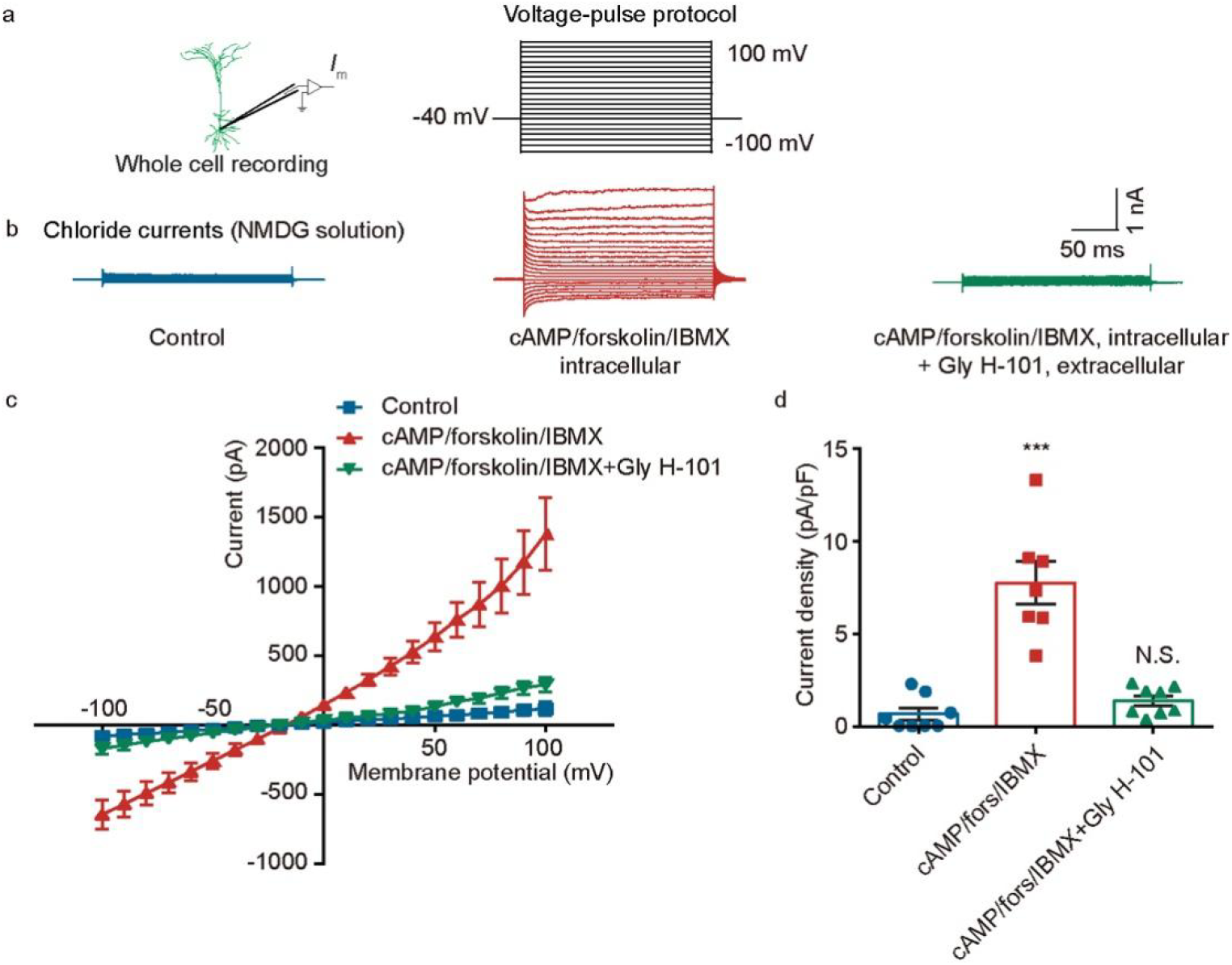
CFTR is functionally active in the deep layer of the cerebral cortex. **a**. Whole-cell stepped voltage-clamp recorded the chloride currents of pyramidal neurons. **b**. N-methyl-D-glucamine (NMDG) solution without Na^+^, K^+^, and Ca^2+^ is utilized for chlorine current recording. CFTR-mediated Cl^-^ currents were induced by the cAMP complex in the pipette solution, which was reduced by the CFTR specific inhibitor Gly H-101 (10 uM). **c**. The current-voltage relationship under various pharmacological circumstances. **d**. the current density under different drug conditions. **d**, Data are mean ±s.e.m. *** *P* < 0.001. n = 8 mice.

Together, the results show that CFTR function is present in the layer V/VI pyramidal neurons of the frontal cortex. This suggests that the activation state of these neurons may cause the opening of CFTR channels, which could impact their excitability and signal transmission.

### CFTR activity affects the excitability of pyramidal neurons in the deep cortex

The results above indicate that the chloride current in pyramidal neurons is weak in the resting state. However, when the intracellular cAMP complex activates CFTR, a larger chloride ion current can be observed. These findings suggest that activating CFTR may alter the resting membrane potential. Using a whole-cell current clamp to record the resting membrane potential of layer V pyramidal neurons in the frontal cortex, we found that the CFTR agonist RP-107 may hyperpolarize the resting membrane potential, which is then recovered after further washing (Figure 3a). Using another CFTR agonist, CBIQ, in the extracellular bath solution or the cAMP complex in the intracellular pipette solution produces similar effects (Figure 3b), which can be reversed by CFTR inhibitors (Figures 3c and 3d). On the other hand, CFTR inhibitors do not significantly alter the resting membrane potential of pyramidal neurons (Figure 3b).

**Figure 3.**
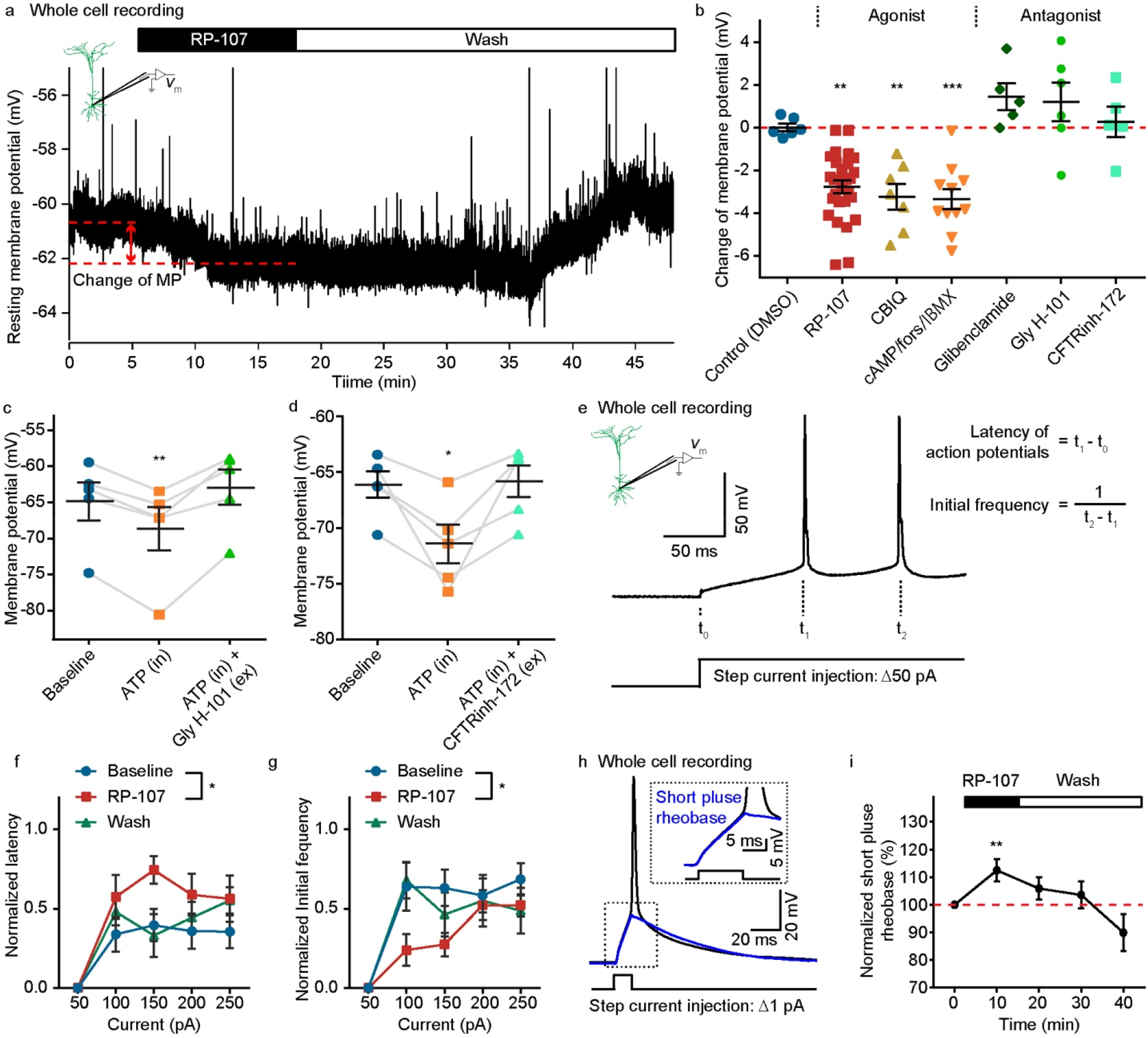
Activating CFTR can hyperpolarize the resting membrane potential of pyramidal neurons and modify the evoked action potential. **a**. Example of whole-cell current-clamp recorded the resting membrane potential of neurons before and after applying the CFTR-specific agonist RP-107. n = 35 mice. **b**. Effect of DMSO, agonists (RP-107 10 uM, CBIQ 10 uM, cAMP complex: cAMP 20 mM/Forskolin 10mM/IBMX 30mM), and antagonists (Glibenclamide 50 uM, Gly H-101 10 uM, CFTRinh-172 10uM) on the resting membrane potential of neurons. n = 6-35 mice. **c d**. A high concentration of intracellular ATP hyperpolarized the resting membrane potential of neurons, which could be blocked by applying the CFTR antagonist (Gly H-101, CFTRinh-172) in the bath solution. n = 5 mice. **e**. Firing action potential of deep pyramidal neurons in response to 800 ms suprathreshold step current injections (ΔI=50 pA) in frontal cortex slices. **f**. Normalized latency of evoked AP with 10 uM RP-107. **g**. Normalized initial firing frequency in the deep frontal cortex with 10 uM RP-107. n = 17 mice. **h**. Inject a gradually increasing short-pulse current to determine the short-pulse rheobase. The blue curve represents the last subliminal current before the evoked action potential. (ΔI=1 pA) **i**, The change of short-pulse rheobase before and after RP-107 treatment. n = 7 mice. **b**,**c**,**d**,**f**,**g**,**i**, Data are mean ±s.e.m. * *P* < 0.05, ** *P* < 0.01, *** *P* < 0.001. n = 6-35 mice.

To further investigate the role of CFTR in regulating neuronal activity, we injected a 50 pA step current (ranging from -250 to 200 pA) into pyramidal neurons and recorded various indicators of evoked action potentials. The results showed that although there was a potential trend in the number of action potentials and input resistance generated by the same stimulus current before and after CFTR activation, it was not statistically significant (Supplementary Figure 6). However, CFTR activation significantly extended the latency of action potentials and decreased the initial frequency (Fig. 3e-g). To determine the effect of CFTR on the evoked action potential threshold of neurons, we injected a positive current into the neuron with a current step of 1 pA and recorded the last stimulus current before the first stable evoked action potential. We found that the threshold current of pyramidal neurons increased after RP-107 treatment and returned to baseline after elution (Fig. 3 h,i).

Overall, neurons exhibited signs of down-regulation of excitability after CFTR activation, such as hyperpolarization of the resting membrane potential and delayed response of evoked action potentials. These findings suggest that the CFTR channel has a negative effect on neuronal excitation.

### Regulating CFTR significantly affects neuronal synaptic transmission

The CFTR protein has the potential to influence neuronal excitability and synaptic transmission. Our experiments, using a whole-cell voltage clamp to record miniature postsynaptic currents in pyramidal neurons, showed that activating the CFTR significantly increased the frequency of mIPSCs (as shown in Figure 4a-e) and slightly decreased the frequency of mEPSCs (as seen in Supplementary Figure 7). On the other hand, inhibiting CFTR channels did not significantly alter synaptic transmission (as depicted in Supplementary Figure 7).

**Figure 4.**
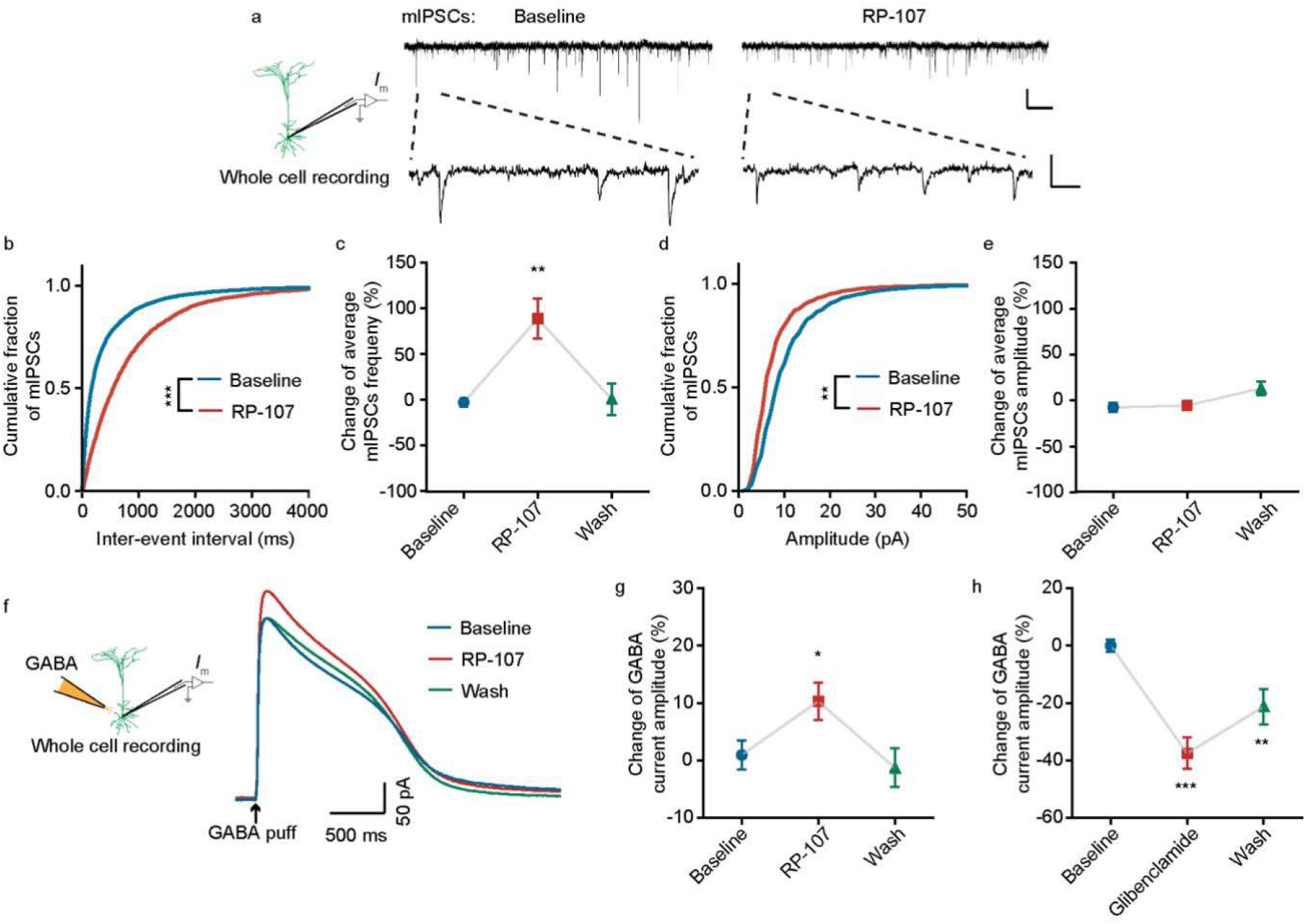
CFTR contributes to inhibitory synaptic transmission. **a**. Example traces of miniature inhibitory postsynaptic currents (mIPSCs) recorded from deep pyramidal neurons of frontal cortex. **b d**. Cumulative fraction analysis of the population inter-event interval and amplitude of mIPSCs with or without RP-107 (10 uM), respectively. **c e**. Mean inter-event interval and amplitude of mIPSCs with or without RP-107, respectively. n = 10 mice. **f**. Example traces of whole-cell currents induced by puffing GABA (1mM) in the deep neurons. **g**. RP-107 increased the amplitude of the GABA-mediated inhibitory current. **h**. Glibenclamide reduced GABA-mediated inhibitory current. n = 8 mice. **b, c, d, g, h**, Data are mean ±s.e.m. * *P* < 0.05, ** *P* < 0.01, *** *P* < 0.001.

To further evaluate the influence of CFTR on inhibitory neurotransmission, we administered 1 mM Gamma aminobutyric acid (GABA), an inhibitory neurotransmitter, directly to the recorded soma. By using the agonist RP-107 (5 mM) in the bath solution, we observed a significant increase in the postsynaptic current mediated by GABA. In contrast, when CFTR channels were inhibited, the peak amplitude of the GABA current was significantly reduced. This drug-induced inhibitory current offset was restored after being exposed to extracellular bath solution (as shown in Figure 4f-h).

In conclusion, the regulation of CFTR activity can impact inhibitory synaptic transmission in neurons, although its effect on excitatory transmission is weaker. These regulatory effects appear to occur presynaptically, which is consistent with the distribution of CFTR on axons.

### Inhibition of CFTR changes the learning process in mice

The results of our electrophysiological study on isolated brain slices suggest that CFTR has a limited impact in the resting state but plays a role in the regulation of excitability following activation. In addition, it affects synaptic transmission in neurons. These findings indicate that CFTR may influence activity-dependent neuronal behavior, such as motor skill learning. To further investigate the effect of CFTR on the learning and memory process of motor skills, we performed a study using a variable-speed wheel. We injected either ACSF or CFTRinh-172 into the deep layer (∼500 μm) of the primary motor cortex on both sides of the mice. After 30 minutes of recovery, the mice were tested for their performance in rotating a rod forward. The memory of this task was then measured again after 20 backward interferences. Our results show that inhibition of CFTR improved the motor learning performance of the mice compared to the ACSF group (Supplementary Figure 8).

The previous research has shown that CFTR is predominantly expressed in the bodies and axons of neurons and may play a role in signal transmission following neuron activation. To further investigate this in vivo, we utilized a two-photon microscope to image the calcium ion indicator GCaMP6s in the heads of living mice that had been fixed in place. Through training on treadmills, we were able to observe the calcium signal in the axon boutons of these neurons (Figure 5a-c). Upon the application of CFTRinh-172 (20 uM), we found that the calcium activity of the axon boutons projected to the imaging area significantly increased (Figure 5d-j). However, there was no notable difference in the backward learning process when CFTR was continuously inhibited (Figure 5k,l).

**Figure 5.**
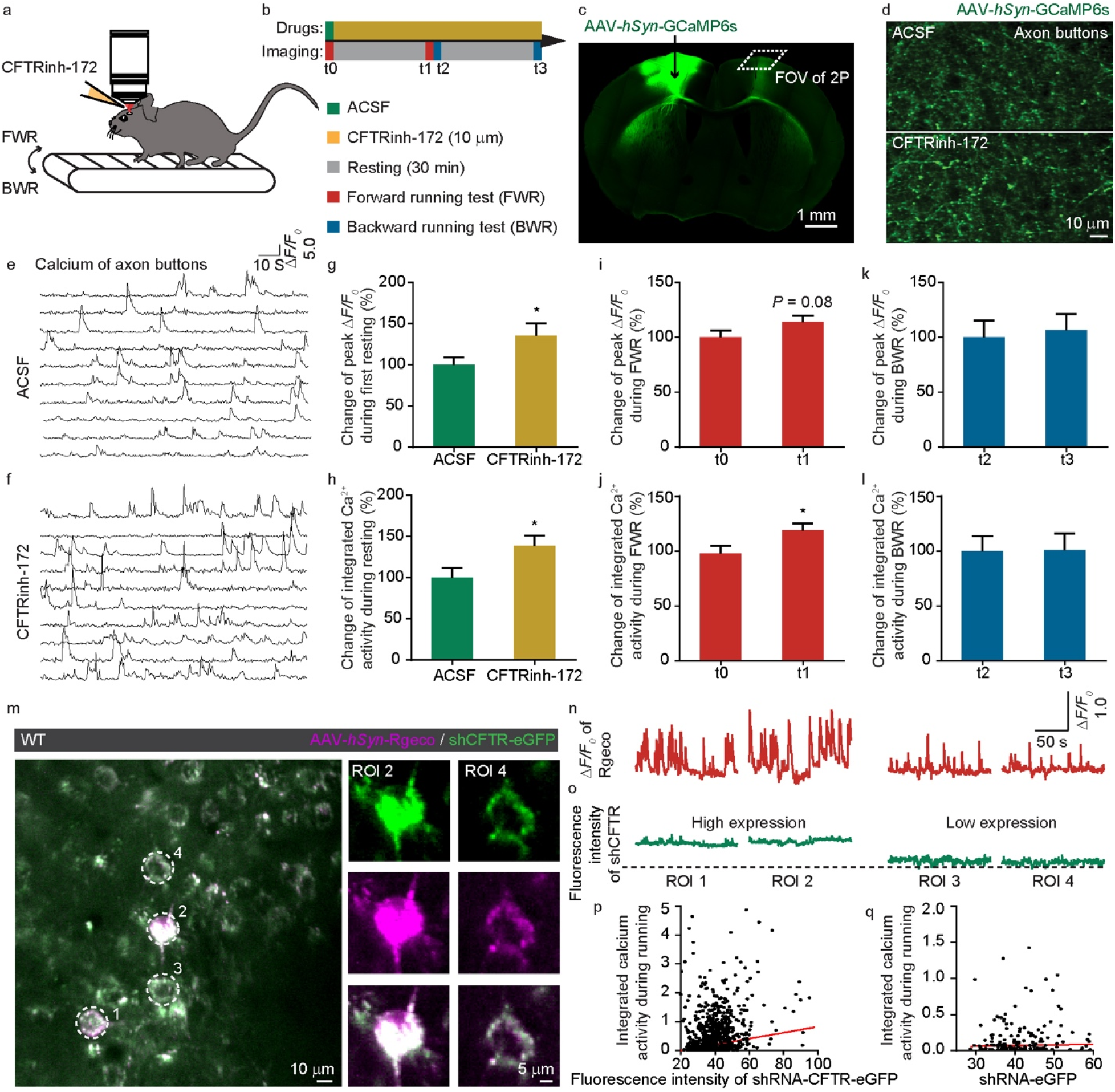
Inhibition of CFTR increases the calcium activity of axon and soma of motor cortex. **a**. Schematic of *in vivo* two-photon calcium imaging of axon boutons or somata while the head-restrained mouse ran forward or backward on a treadmill (FWR or BWR). **b**. The experimental design to detect inhibiting CFTR on the learning process. **c**. We injected GCaMP6s virus into layer V (∼500 um) of the primary motor cortex of the mouse and imaged the axon boutons projected to the contralateral cortex. **d**. Calcium fluorescence of projected axon boutons with ACSF or CFTRinh-172 (20 uM). **e f**. Representative running-induced calcium traces of projected axon boutons with ACSF or CFTRinh-172 bath solution. **g**. Peak Δ*F/F*_*0*_ of calcium transients increased after CFTRinh-172 application in the first resting period. **h**. Integrated calcium activity increased after CFTRinh -172 application in the first resting period. **i**. Peak Δ*F/F*_*0*_ of FWR-induced calcium spikes had an increasing trend after CFTRinh-172 treatment. **j**. FWR-induced integrated calcium activity increased after CFTRinh-172 application. **k l**, Continuous bathing with CFTRinh-172 had no effect on the peak Δ*F/F*_*0*_ or integrated calcium activity during BWR learning. n = 33 FOVs, 13 mice **m**. Representative image of Rgeco (magenta) labelled calcium signals with eGFP (green) labelled sh-CFTR lentivirus. **n o**, The Δ*F/F*_*0*_ of calcium activity with different expressions of eGFP. On the left, cells with strong expression of the green lentivirus have strong calcium activity. The right side indicates cells with weaker green fluorescence and lower calcium activity. **p**. Correlation analysis between the fluorescence value of the green sh-CFTR-eGFP lentivirus and the integrated activity of red calcium. **q**. The control group, sh-eGFP lentivirus, showed no correlation. n = 3 mice. **g, h, j**, Data are mean ±s.e.m. * *P* < 0.05.

To confirm the role of the CFTR channel in the bodies of living animals, we used a short hairpin RNA lentivirus targeting the cftr gene to reduce its expression in neurons. This lentivirus, along with AAV-Syn-Rgeco, was injected into the primary motor cortex of mice. Upon double infection, we observed that the eGFP fluorescence intensity of neurons infected by the shRNA virus was varied (Figure 5m,o). Additionally, the red calcium activity during the treadmill learning process w xcas also different (Figure 5n). We found a positive correlation between the fluorescence intensity of the green shRNA lentivirus and red calcium activity during motor learning (Figure 5p). However, this correlation was not present in the control virus that only expressed shRNA-eGFP (Figure 5q). Overall, our findings suggest that inhibiting or reducing the CFTR channel in the primary motor cortex of mice can alter the motor learning process. We also observed enhanced somatic and axonal calcium activity in the motor cortex of living animals.

## 3. Discussion

Through extensive studies on the CFTR channel in epithelial tissues, it can be speculated that CFTR in the nervous system plays a crucial role in three main aspects. The first hypothesis suggests that CFTR can directly permeate chloride ions, thereby maintaining neuronal chloride balance, cell homeostasis, and regulating neuronal excitability and synaptic transmission. In this study, we focused mainly on hypothesis 1. We scanned the distribution of CFTR in the brains of adult rats and confirmed the CFTR protein’s cell type in brain slices using immunological double labelling. Our findings revealed that CFTR was mainly distributed in the deep pyramidal somata and superficial axons of the cerebral cortex. We employed the patch-clamp technique to confirm the existence of the chloride current activated by cAMP in the deep pyramidal neurons of the cerebral cortex, which could be blocked by CFTR-specific inhibitors. Furthermore, activation of the CFTR channel could hyperpolarize the resting membrane potential, delay the evoked action potential, and increase the presynaptic effect of inhibitory synaptic transmission. Inhibiting or reducing the CFTR channel could increase the neuronal calcium activity of the soma and contralateral axon in vivo and even improve the motor learning process.

In this study, we aimed to systematically investigate the function of CFTR using a variety of methods. We obtained some consistent results, including the distribution of CFTR on axons matching the presynaptic effect of CFTR electrophysiologically and calcium activity changes in the axon boutons projected to the contralateral cortex observed in in vivo experiments. However, some results did not seem to be the same, such as the insignificant direct inhibition of CFTR in isolated brain slices, but it could play a blocking role when CFTR was activated. For in vivo experiments, inhibiting CFTR could directly change the calcium activity of neurons. One possible explanation for this inconsistency is that the nerve projections from various brain regions were severed during the preparation of the brain slice. As a result, the activity of nerve cells is significantly higher in living animals with complete neural circuits than in isolated tissues. Therefore, inhibiting CFTR may have an observable effect on neuronal homeostasis and chloride balance. This speculation, however, needs to be confirmed through chlorine imaging in living animals.

It is worth noting that we studied the function of CFTR in mature rodent neurons, and the neuronal chlorine balance is reversed with the maturation of various chlorine channels and transporters, such as GABAR, KCC2, NKCC1, etc. This developmental process suggests that CFTR functions may differ in the early stages of development and maturation. Further research is needed to determine whether CFTR has a synergistic effect or functional changes with other chloride channels or transporters^44–46^.

Hypothesis 2 proposes that CFTR functions as a small molecule channel in close proximity to the cellular membrane, regulating the release of ATP, GSH, and enkephalin^47–49^. Evidence from studies on SD rats demonstrates that glutamate in the spinal cord increases intracellular calcium concentration, resulting in the release of ATP by microglia. However, the mutation of CFTR inhibits this release. Moreover, CFTR also impedes the release of ATP mediated by adrenaline beta-3 receptors on sensory neurons in the dorsal root ganglion.

In addition, hypothesis 3 posits that CFTR may be linked to cytoskeletal proteins through the PDZ domain under the membrane, enabling the regulation of other receptors or channels^50,51^. Upon activation of the adrenaline beta-2 receptor, the Gs signaling pathway triggers an increase in the cell’s local cAMP concentration, leading to PKA phosphorylation of the R domain of CFTR and alteration of the flow of local chloride ions. Further research is necessary to determine if CFTR exhibits similar effects in the brain as proposed in hypotheses 2 and 3.

Maintaining a stable and functional central nervous system necessitates a balance between excitation and inhibition. The occurrence of brain disorders and damage is likely with excessive excitation or inhibition. The regulation of neural network activity, which is critical for maintaining anion balance, is an essential role of chloride channels. Nonetheless, the study of chloride channels lags behind that of other cation families. Our study thoroughly examined the function of the cystic fibrosis transmembrane regulator (CFTR) in the cerebral cortex, and it is of great theoretical significance in clarifying the mechanism of chloride ion channels in the central nervous system.

## 4. Materials and Methods

### Animal preparation

Sprague Dawley (SD) rats and C57/BL6 mice were used in this study. SD rat breeders were purchased from the Experimental Animal Center of Nanchang University, and C57/BL6 mice were purchased from the Jackson Lab Animal Center in the United States. Our laboratory reproduced the subsequent use of experimental animals. 2-3 adult rats or five mice are kept in each cage during the breeding process. The experimental animals were given sufficient feed and water to maintain a 12/12-hour photoperiod. All operations involving animal experiments follow the “Guidelines for the Care and Use of Laboratory Animals” promulgated by the National Institutes of Health (NIH). Minimize the number of animals used in the experimental design process and minimize the pain of animals during the surgery process.

### Immunohistochemistry (IHC)

SD rats aged seven weeks (P49) were perfused and fixed with 4% paraformaldehyde. Cut the rat brain tissue in a coronal and sagittal pattern with a thickness of 30 μm on a cryostat. The brain sections were washed thoroughly with PBS and incubated with CFTR primary antibody for 12 hours at 4°C. Four CFTR antibodies were used in this study: Mouse anti-CFTR (Thermofisher, M3A7, 1:100), Rabbit anti-CFTR (Abcam, ab117447, 1:200), Mouse anti-CFTR (Abcam, CF3, 1:100)), Rabbit anti-CFTR (Alomone Labs, ACL-006, 1:100).The verification of antibodies specificity is shown in Supplementary Figure 1. After the primary antibody reaction, the section was incubated with AlexaFluor-conjugated secondary antibody (488 or 594) for 2 hours at room temperature and washed. To visualize the nucleus DAPI staining (Invitrogen) was performed. For other markers, use the following markers: Mouse anti-CaMK II, (Abcam, ab52476, 1:350), Mouse anti-GAD65/67, (Abcam, ab11070, 1:1000), Mouse anti-NF200, (Abcam, ab82259, 1:2000), Mouse anti-MAP 2, (Abcam, AB24640, 1:1000), Mouse anti-Beta-tubulin III, (Abcam, AB18207, 1:1000). After antibody staining, laser confocal observation was performed on those brain sections.

### Cell culture and Immunocytochemistry (ICC)

The culture of cerebral cortex neurons is done as previously described. Briefly, the cerebral cortex from E18-19 embryos should be collected and separated in ice-cold HBSS (Gibco). After 20 minutes of digestion in 0.25% trypsin at 37°C. Resuspend the dissociated cells in a plating medium (DMEM supplemented with 10% FBS) and distribute them at a density of 3E4 or 6E4 per well on poly-L-lysine-coated 8 mm coverslips (Fisher). The cells were cultured for 4 hours before being switched to a maintenance medium [Neural Basal Medium (Gibco) with 2% B-27 supplement (Gibco), 1% GlutaMax (Gibco), and 1% Penicillin/Streptomycin (Gemini)]. The neurons are kept at 37 °C in 5% CO2, with half of the medium replaced every two days.

Morphological studies were performed 21 days later. The neurons were fixed with 4% paraformaldehyde for 30 minutes at 4°C. The cell slides were blocked with PBS containing 5% goat serum at 4°C, and permeabilized with 0.1% Triton X-100 at room temperature for 5 minutes. These cells were treated with an anti-CFTR antibody and neuron-specific tubulin III. Following the reaction of the primary antibody, the secondary antibody (488 or 594) conjugated with AlexaFluor is incubated for 2 hours at room temperature before being washed. DAPI staining (Invitrogen) was used to visualize the nucleus.

### Patch clamp recordings

SD rats (5-week-old males) were anaesthetized with sodium pentobarbital (Sigma, 40 mg/kg, intraperitoneal injection). The brain is quickly removed and cooled in ice-cold modified artificial cerebrospinal fluid (ACSF), which contains (in millimoles): 250 glycerol, 2 KCl, 10 MgSO_4_, 0.2 CaCl_2_, 1.3 NaH_2_PO_4_, 26 NaHCO_3_, and 10 glucoses. Use VT-1200S vibratome (Leica) to cut coronal sections (300 μm) in ice-cold modified ACSF, and then transfer to conventional ACSF (mmol, 126 NaCl, 3 KCl, 1 MgSO_4_, 2 CaCl_2_, 1.25 NaH_2_PO_4_, 26 NaHCO_3_, and 10 Glucose). Keep it at 30 °C for 30-60 minutes, and then keep it at room temperature (25 ± 1 °C) before recording. All solutions were saturated with 95% O_2_/5% CO_2_ (vol/vol). The slices were placed in the recording chamber and perfused with ACSF (2 ml/min) at 25°C. We used an upright microscope equipped with a 40x water immersion lens (Axioskop 2 Plus, Zeiss) and an infrared-sensitive CCD camera (C2400-75, Hamamatsu) to observe the nerve cells in the cerebral cortex with an infrared optical system. Triangular cells, more than 500um away from the cerebral cortex, were selected for recording. The pipette was pulled by a micropipette puller (P-97, Sutter) with a resistance of 3-5 MΩ and filled with different electrode pipette solutions. We used MultiClamp 200B amplifier and 1440A digitizer (Molecular Devices) to record whole-cell patch-clamp configuration. The series resistance was controlled below 20 MΩ and was not compensated. In each whole-cell patch-clamp experiment recording, the first five minutes were used as the process of intracellular and electrode balancing. After the series impedance was stable, the subsequent drug experiment was carried out. The data were filtered at 1 kHz and sampled at 10 kHz. All experiments were performed at room temperature (25 °C).

For the chloride current recording, we clamped the cells at -40 mV. The adjusted N-methyl-D-glucamine (NMDG) solution was used to eliminate the influence of cations. The pipette (intracellular) solution contained (in mM): 140 N-methyl-D-glucamine, 40 HCl, 100 L-glutamic acid, 0.2 CaCl_2_, 2 MgCl_2_, 1 EGTA, 10 HEPES, 2 ATP-Mg, pH 7.3, with HCl. The bath (extracellular) solution contained (in mM): 140 N-methyl-D-glucamine, 10 HEPES, 1 CaCl_2_,1 MgCl_2_. For the recording of membrane potential, action potential and excitatory synaptic transmission, the cells were clamped at -70 mV with the following pipette intracellular solutions (in mM):150 K-Gluarate, 0.4 EGTA, 8 NaCl, 10 HEPES, 2 ATP-Mg, 0.1 GTP-Na, 10 Na-Phosphocrateine. For inhibitory synaptic transmission recording, the cells were clamped at -40 mV with the following pipette intracellular solutions (in mM): 70 K-Gluarate, 70 KCl, 20 HEPES, 0.5 CaCl_2_, 5 EGTA, 5 ATP-Mg.

### Drug application

The drugs were applied at 5 min for all pharmacological experiments, and the elution started at 17 min. The total recording time was approximately 50-60 min. The drugs used in the experiment are as follows: CFTR agonist: RP-107 10 uM, CBIQ 10 uM, cAMP Complex: cAMP 20 mM/Forskolin 10 mM/IBMX, 30 mM. CFTR antagonist: CFTRinh-172 10-20 uM, Gly H-101, 5 uM, Glibenclamide 50-100 uM. During cell-attached recorded chloride current, Picrotoxin 100 uM, Tetrodotoxin 1uM were introduced to the extracellular bathing solution.1 mM GABA solution was used to record the chloride current mediated by inhibitory transmitters.

RP-107, CBIQ, cAMP, Forskolin, IBMX, CFTRinh-172, Gly H-101, Glibenclamide, Picrotoxin, GABA, ATPMg, GTPNa^3+^, and HEPES were purchased from sigma chemical company (Sigma, St Louis, MO, USA). Tetrodotoxin was obtained from the research institute of aquatic products, Hebei province, China.

### Rotating rod experiment

To study the effect of CFTR in the cortex on motor skill performance, we used a uniform acceleration rotating rod instrument for training and observation. According to Yang’s method^52^, in brief: during forwarding rotation, the speed of the rotating rod is set to increase from 0 to 100 rpm within 3 minutes. During training, the animal needs to run with a rotating rod. We recorded the time when the animal fell. During reverse rotation, the maximum speed was set to increase to 30 rpm within 3 minutes.

The mice in the control and experimental groups had their heads mounted with a self-made head bar one day before the test. Slowly injected CFTRinh-172 (20 uM) or ACSF into the primary motor cortex on both sides of the mouse through a pressure delivery device. The injection pressure was 30 psi, the injection frequency was 0.1 Hz, and each injection lasted 20 ms. A total of 500 uL was injected into each brain area for 5 min. After the injection, stay for 2 minutes to ensure that the drug was completely absorbed. Pull out the electrode, seal the injection hole with medical silica gel, and recover for 30 minutes for rotate training.

### Two photon imaging

The experimental animals were deeply anaesthetized with ketamine/xylazine the day before imaging. Mounted a self-made head fixation device on the head and exposed the target area. Polish the top of the imaging area on the day of imaging, remove the skull and install 0# glass. Use glue to fix the glass with the skull and leave a hole where the drug can penetrate. The neurons were then visualized using a 25X water objective lens. We used the Burker two-photon system with a 920 nm wavelength laser. The max laser power was 300 mW pass lens.

The motor learning process uses treadmill training regarding xx. Ensure that the two front legs could move freely when the head was fixed. During training, a 10 V DC power supply drove the treadmill motor (Dayton, model 2L010). Each running training lasts 60 seconds. After each run, turn off the treadmill and start the next test after a 90-second rest period. At 15 s before each run, two-photon imaging was started, and the imaging time lasted for 90 s (that is, including 15 s resting period, 60 s running period and 15 s post-running resting period).

### ShRNA expression

To interfere with the expression of *cftr*, the Vigene bioscience-packaged lentivirus (5’-CCGGGCTGAAAGCAGGTGGGATTCTCAAGAGAAATCCCACCTGCTTTCAGCTTTTTTG-3’^53^) was used for this study. Calcium imaging was performed using the AAV-Syn-Rgeco virus. The AAV virus was injected into the mouse’s primary motor cortex at a depth of 500 microns using stereotaxic. One week after Rgeco virus expression, the Sh-CFTR-eGFP lentivirus was injected. The mice were allowed to express for 7-10 days after receiving the two viruses for two-photon imaging.

### Data analysis

#### Patch-clamp signal processing

The statistical results of all patch clamps are calculated using *clamp fit 9*.*0* software, including peak potential amplitude, time, average voltage intensity, current-voltage curve, etc. For mI/EPSCs data, use *clamp fit 9*.*0* for conversion and import the data into the *mini-Analysis program* for measurement. The main measurement parameters include the amplitude of the miniature current, the time interval, and the curve’s area. Neurons whose resistance fluctuates more than 20% are not counted.

#### Two-photon data processing

*Image J* software is primarily used to analyze images in the two-photon imaging process. We analyze the changes in fluorescence intensity and calcium ion signal.

For the analysis of calcium signal, each mouse selects 3-5 fields of the primary motor cortex, each area has a width of 235 µm (Zoom3) or 142 µm (Zoom5), and a resolution of 1024×1024 pixels. The frame rate is 0.72 Hz. The region of interest (Region of interests, ROI) is chosen with the selection tool of Image J software, and the average fluorescence value in the ROI area of each frame is counted. Use Δ*F/F*_*0*_ to indicate the specific change in the fluorescence value. Select the minimum 10% of each cell as F0 during the resting state. In the running condition, the average value of the last three seconds is *F*_*0*_. The peak amplitude is defined as the maximum value of each calcium transient. The area under the peak calcium traces is divided by the time to obtain the integrated calcium activity^54^.

### Statistical analysis

All data in this experiment are expressed as mean ± standard error. All statistical data are firstly tested by normal distribution analysis (Kolmogorov-Smirnov test) to determine whether the experimental data can be tested with parameters. When the distribution of the experimental data conforms to the normal distribution, the two-tailed Student t-test is selected to compare the two groups of data, and the two-way ANOVA test is selected for the comparison between multiple groups. The detection process selects paired or unpaired detection according to the specific conditions of the experiment. For experimental data that does not conform to the normal distribution, non-parametric detection is selected. All significant differences are represented by *, where * *P*<0.05, ** *P*<0.01, *** *P*<0.001. Data greater than 0.05 are considered to have no statistically significant difference. All statistical calculations are done using *Graphpad Prism*.

## Acknowledgments

We thank all the members in the Li Laboratory and the Gan Laboratory for their constructive discussion on this project. Z.C. thanks Dr. Bingxing Pan, Dr. Xuhui Zeng, and Dr. Hong Xue (Nanchang University) for their helpful comments and discussion.

## Author contributions

B.L., and H.C.C. initiated the project. B.L., and W.G. supervised the research. Z.C. and X.S. designed and carried out the biology experiment. Z.C., X.S., and W.X. performed data analysis and statistics. L.F. taught patch clamp technique. Z.C., and B.L. contributed to the writing of the manuscript. All authors read and approved the manuscript.

## Data and codes availability

All the data and codes described in this study can be obtained from the authors upon reasonable request.

## Supplemental document

See Supplement information 1 for supporting content.

## References

1. Riordan, J. R. et al. Identification of the cystic fibrosis gene: Cloning and characterization of complementary DNA. Science (1979) 245, 1066–1073 1989.

2. M. Rowe, steven□ M.D., Stacey Miller, B.S., and Eric J. Sorscher, M. D. Cystic fibrosis Cystic fibrosis. Journal of Cystic Fibrosis 21, 1–378 2022.

3. Greger, R. et al. Cystic fibrosis and CFTR. Pflugers Arch 443, 3–7 2001.

4. Cant, N., Pollock, N. & Ford, R. C. CFTR structure and cystic fibrosis. International Journal of Biochemistry and Cell Biology 52, 15–25 2014.

5. Rommens, J. M. et al. Cystic Identification of the Chromosome Walking Fibrosis Gene_J: Jumping and. Adv Sci 245, 1059–1065 2009.

6. Quinton, P. M. Cystic fibrosis: lessons from the sweat gland. Physiology (Bethesda) 22, 212–225 2007.

7. Blackman, S. M. et al. Relative contribution of genetic and nongenetic modifiers to intestinal obstruction in cystic fibrosis. Gastroenterology 131, 1030–1039 2006.

8. Hardin, D. S. GH improves growth and clinical status in children with cystic fibrosis -- a review of published studies. Eur J Endocrinol 151 Suppl 1, 2004.

9. Flume, P. A. et al. Cystic Fibrosis Pulmonary Guidelines. 10.1164/rccm.201002-0157OC 182, 298–306 2012.

10. Cohn, J. A. et al. Relation between mutations of the cystic fibrosis gene and idiopathic pancreatitis. N Engl J Med 339, 653–658 1998.

11. Malfroot, A. & Dab, I. New insights on gastro-oesophageal reflux in cystic fibrosis by longitudinal follow up. Arch Dis Child 66, 1339–1345 1991.

12. Kalayoglu, M., Sieber, W. K., Rodnan, J. B. & Kiesewetter, W. B. Meconium ileus: A critical review of treatment and eventual prognosis. J Pediatr Surg 6, 290–300 1971.

13. Tremellen, K. Oxidative stress and male infertility—a clinical perspective. Hum Reprod Update 14, 243–258 2008.

14. McCray, P. B. et al. Expression of CFTR and presence of cAMP-mediated fluid secretion in human fetal lung. Am J Physiol 262, 1992.

15. Yoshimura, K. et al. Expression of the human cystic fibrosis transmembrane conductance regulator gene in the mouse lung after in vivo intratracheal plasmid-mediated gene transfer. Nucleic Acids Res 20, 3233–3240 1992.

16. Dray-Charier, N. et al. Expression of delta F508 cystic fibrosis transmembrane conductance regulator protein and related chloride transport properties in the gallbladder epithelium from cystic fibrosis patients. Hepatology 29, 1624–1634 1999.

17. Elgavish, A. High intracellular pH in CFPAC: a pancreas cell line from a patient with cystic fibrosis is lowered by retrovirus-mediated CFTR gene transfer. Biochem Biophys Res Commun 180, 342–348 1991.

18. Fonknechten, N. et al. Skipping of exon 9 in CFTR mRNA of human adult and fetal pancreas from non-CF individuals. Hum Mol Genet 2, 2141–2142 1993.

19. Cohn, J. A., Melhus, O., Page, L. J., Dittrich, K. L. & Vigna, S. R. CFTR: development of high-affinity antibodies and localization in sweat gland. Biochem Biophys Res Commun 181, 36–43 1991.

20. Morales, M. M. et al. Both the wild type and a functional isoform of CFTR are expressed in kidney. Am J Physiol 270, 1996.

21. Meschede, D., Eigel, A., Horst, J. & Nieschlag, E. Compound heterozygosity for the delta F508 and F508C cystic fibrosis transmembrane conductance regulator (CFTR) mutations in a patient with congenital bilateral aplasia of the vas deferens. Am J Hum Genet 53, 292 1993.

22. Kunzelmann, K. Introduction to section V: assessment of CFTR function. Methods Mol Biol 741, 407–418 2011.

23. Reznikov, L. R. et al. CFTR-deficient pigs display peripheral nervous system defects at birth. Proc Natl Acad Sci U S A 110, 3083–3088 2013.

24. Yeh, K. M. et al. Cystic fibrosis transmembrane conductance regulator modulates enteric cholinergic activities and is abnormally expressed in the enteric ganglia of patients with slow transit constipation. J Gastroenterol 54, 994–1006 2019.

25. Zhang, Y. P. et al. CFTR prevents neuronal apoptosis following cerebral ischemia reperfusion via regulating mitochondrial oxidative stress. J Mol Med (Berl) 96, 611–620 2018.

26. Pan, P., Guo, Y. & Gu, J. Expression of cystic fibrosis transmembrane conductance regulator in ganglion cells of the hearts. Neurosci Lett 441, 35–38 2008.

27. Niu, N., Zhang, J., Guo, Y., Yang, C. & Gu, J. Cystic fibrosis transmembrane conductance regulator expression in human spinal and sympathetic ganglia. Lab Invest 89, 636–644 2009.

28. Su, M., Guo, Y., Zhao, Y., Korteweg, C. & Gu, J. Expression of cystic fibrosis transmembrane conductance regulator in paracervical ganglia. Biochem Cell Biol 88, 747–755 2010.

29. Mulberg, A. E. et al. Cystic fibrosis transmembrane conductance regulator protein expression in brain. Neuroreport 5, 1684–1688 1994.

30. Marcorelles, P. et al. Cystic Fibrosis Transmembrane Conductance Regulator Protein (CFTR) Expression in the Developing Human Brain: Comparative Immunohistochemical Study between Patients with Normal and Mutated CFTR. Journal of Histochemistry and Cytochemistry 62, 791–801 2014.

31. Hincke, M. T., Nairn, A. C. & Staines, W. A. Cystic fibrosis transmembrane conductance regulator is found within brain ventricular epithelium and choroid plexus. J Neurochem 64, 1662–1668 1995.

32. Pfister, S. et al. Novel role of cystic fibrosis transmembrane conductance regulator in maintaining adult mouse olfactory neuronal homeostasis. J Comp Neurol 523, 406–430 2015.

33. Morales, F. R., Silveira, V., Damián, A., Higgie, R. & Pose, I. The possible additional role of the cystic fibrosis transmembrane regulator to motoneuron inhibition produced by glycine effects. Neuroscience 177, 138–147 2011.

34. Ostroumov, K., Grandolfo, M. & Nistri, A. The effects induced by the sulphonylurea glibenclamide on the neonatal rat spinal cord indicate a novel mechanism to control neuronal excitability and inhibitory neurotransmission. Br J Pharmacol 150, 47–57 2007.

35. Guo, Y., Su, M., McNutt, M. A. & Gu, J. Expression and Distribution of Cystic Fibrosis Transmembrane Conductance Regulator in Neurons of the Human Brain. Journal of Histochemistry and Cytochemistry 57, 1113 2009.

36. Goldstein, A. B. et al. Cystic Fibrosis Patients With and Without Central Nervous System Complications Following Lung Transplantation. Pediatr Pulmonol 30, 203–206 2000.

37. Vaughn, B. V. et al. Seizures in Lung Transplant Recipients. Epilepsia 37, 1175–1179 1996.

38. Gentzsch, M., Choudhury, A., Chang, X. B., Pagano, R. E. & Riordan, J. R. Misassembled mutant DeltaF508 CFTR in the distal secretory pathway alters cellular lipid trafficking. J Cell Sci 120, 447–455 2007.

39. Eworuke, E., Zeng, Q. Y. L. & Winterstein, A. G. Clinical and Sociodemographic Factors Associated With Attention-Deficit/Hyperactivity Disorder in Patients With Cystic Fibrosis. Psychosomatics 56, 495–503 2015.

40. Georgiopoulos, A. M. & Hua, L. L. The Diagnosis and Treatment of Attention Deficit-Hyperactivity Disorder in Children and Adolescents with Cystic Fibrosis: A Retrospective Study. Psychosomatics 52, 160–166 2011.

41. Ostroumov, K. A new stochastic tridimensional model of neonatal rat spinal motoneuron for investigating compartmentalization of neuronal conductances and their influence on firing. J Neurosci Methods 163, 362–372 2007.

42. Reznikov, L. R. Cystic Fibrosis and the Nervous System. Chest 151, 1147 2017.

43. Wu, Y. et al. CFTR Modulates Hypothalamic Neuron Excitability to Maintain Female Cycle. Int J Mol Sci 24, 12572 2023.

44. Wilke, B. U., Kummer, K. K., Leitner, M. G. & Kress, M. Chloride – The Underrated Ion in Nociceptors. Front Neurosci 14, 520280 2020.

45. Henao Romero, N. & 0000-0002-4878-679X. THE ROLE OF THE CYSTIC FIBROSIS TRANSMEMBRANE REGULATOR (CFTR) IN CHLORIDE HOMEOSTASIS AND EXCITABILITY IN SWINE SENSORY NEURONS. 2021.

46. Krishnan, V., Maddox, J. W., Rodriguez, T. & Gleason, E. A role for the cystic fibrosis transmembrane conductance regulator in the nitric oxide-dependent release of Cl-from acidic organelles in amacrine cells. J Neurophysiol 118, 2842–2852 2017.

47. Guo, J. L., Kalous, A., Werry, E. L. & Bennett, M. R. Purine release from spinal cord microglia after elevation of calcium by glutamate. Mol Pharmacol 70, 851–859 2006.

48. Kanno, T., Yaguchi, T. & Nishizaki, T. Noradrenaline stimulates ATP release from DRG neurons by targeting beta(3) adrenoceptors as a factor of neuropathic pain. J Cell Physiol 224, 345–351 2010.

49. Chalmers, J. A., Jang, J. J. & Belsham, D. D. Glucose sensing mechanisms in hypothalamic cell models: glucose inhibition of AgRP synthesis and secretion. Mol Cell Endocrinol 382, 262–270 2014.

50. Su, X. et al. Interregulation of proton-gated Na+ channel 3 and cystic fibrosis transmembrane conductance regulator. Journal of Biological Chemistry 281, 36960–36968 2006.

51. Ji, H. L. et al. Up-regulation of acid-gated Na(+) channels (ASICs) by cystic fibrosis transmembrane conductance regulator co-expression in Xenopus oocytes. J Biol Chem 277, 8395–8405 2002.

52. Yang, G. et al. Sleep promotes branch-specific formation of dendritic spines after learning. Science 344, 1173–1178 2014.

53. Zhao, L. et al. CFTR deficiency aggravates Ang II induced vasoconstriction and hypertension by regulating Ca2+ influx and RhoA/Rock pathway in VSMCs. Front Biosci (Landmark Ed) 26, 1396–1410 2021.

54. Cheng, Z. et al. Probing neuronal functions with precise and targeted laser ablation in the living cortex. Optica 8, 1559 2021.

