## Supplemental information for "Cystic fibrosis transmembrane conductance regulator (CFTR) downregulates the excitability of deep pyramidal neurons in the rodent cortex"

Zongyue Cheng^1,2,3^, Xuan Sun^1^, Wang Xing^1^, Fei Luo^1^, Hsiao Chang Chan^4^,
Wenbiao Gan^5^, Baoming Li^1*^

1 Institute of Life Science and School of Life Science, Nanchang University, Nanchang 330031, China.

2 Skirball Institute, Department of Neuroscience and Physiology, Department of Anesthesiology, New York University School of Medicine, New York, NY 10016, USA

3 School of Electrical and Computer Engineering, Purdue University, West Lafayette, IN 47907, USA

4 Epithelial Cell Biology Research Center, School of Biomedical Sciences, Faculty of Medicine, The Chinese University of Hong Kong, Shatin, Hong Kong, China.

5 Institute for Neurological Diseases, Shenzhen Bay Laboratory, Shenzhen, 518132, China.

**This file includes Supplementary Figure 1-8.**

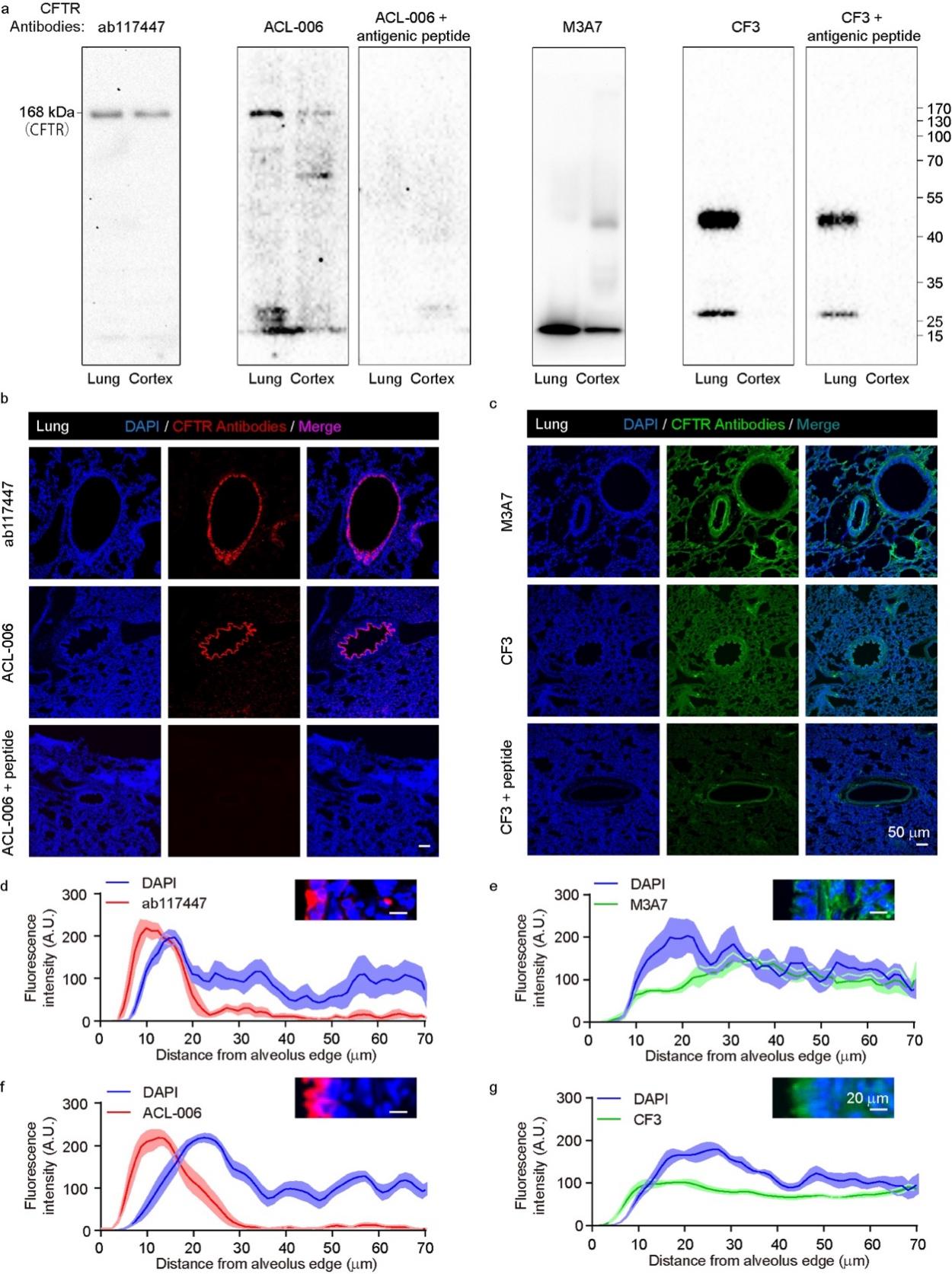

Supplementary Figure 1 | Specificity verification of the four CFTR antibodies. a. Western blotting verified the target band positions of the four CFTR antibodies and the blocking effect of antigen peptides. The size of the blot bands of antibody ab117447 and antibody ACL-006 accorded with the theoretical size of CFTR. The blotting of pre-mixed monoclonal antibody ACL-006 and antigen peptide group was negative. Antibodies M3A7 and CF3 did not produce the expected results. b c. CFTR antibody staining in positive lung tissue. d-g. Distribution of CFTR antibodies on the edge of alveolar tissue. Antibody ab117447 and antibody ACL-006 were mainly distributed in lung epithelial cells, which conformed to the distribution pattern of CFTR, while the distribution of antibodies M3A7 and CF3 was more diffuse.

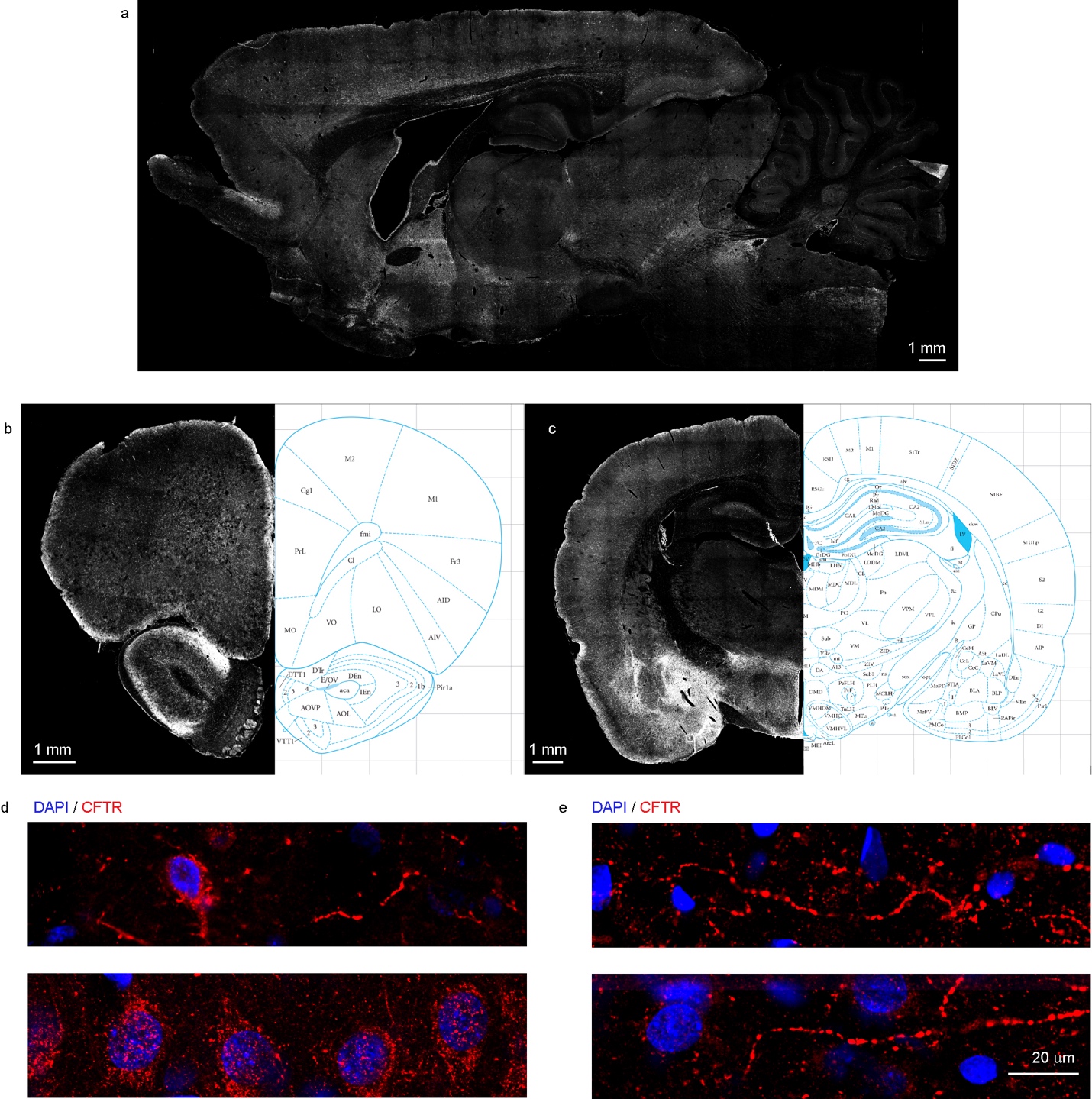

Supplementary Figure 2 | The distribution of CFTR in the rat brain. a. Sagittal distribution of CFTR in the whole rat brain. b. Coronal distribution of CFTR in frontal lobe slices. c. Coronal distribution of CFTR in hippocampus and thalamus. d. CFTR presented cell body-like distribution in the deep cerebral cortex and hippocampus. e. CFTR showed the fibrous distribution in the superficial cerebral cortex, hypothalamus, and midbrain.

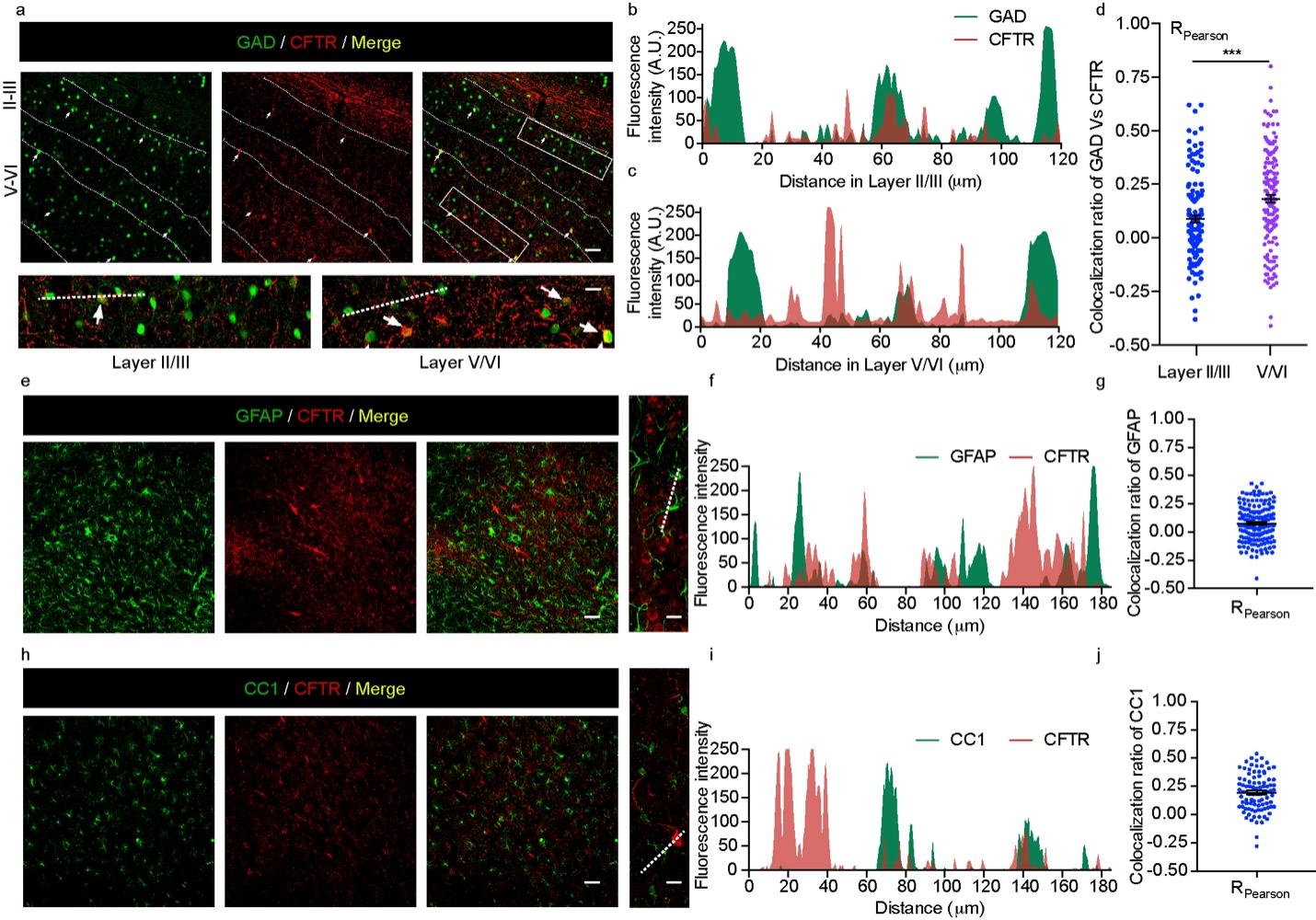

Supplementary Figure 3 | CFTR expression in nerve cells. a-d. CFTR expression in interneurons. e-g. CFTR expression in astrocytes. h-j. CFTR expression in oligodendrocytes.

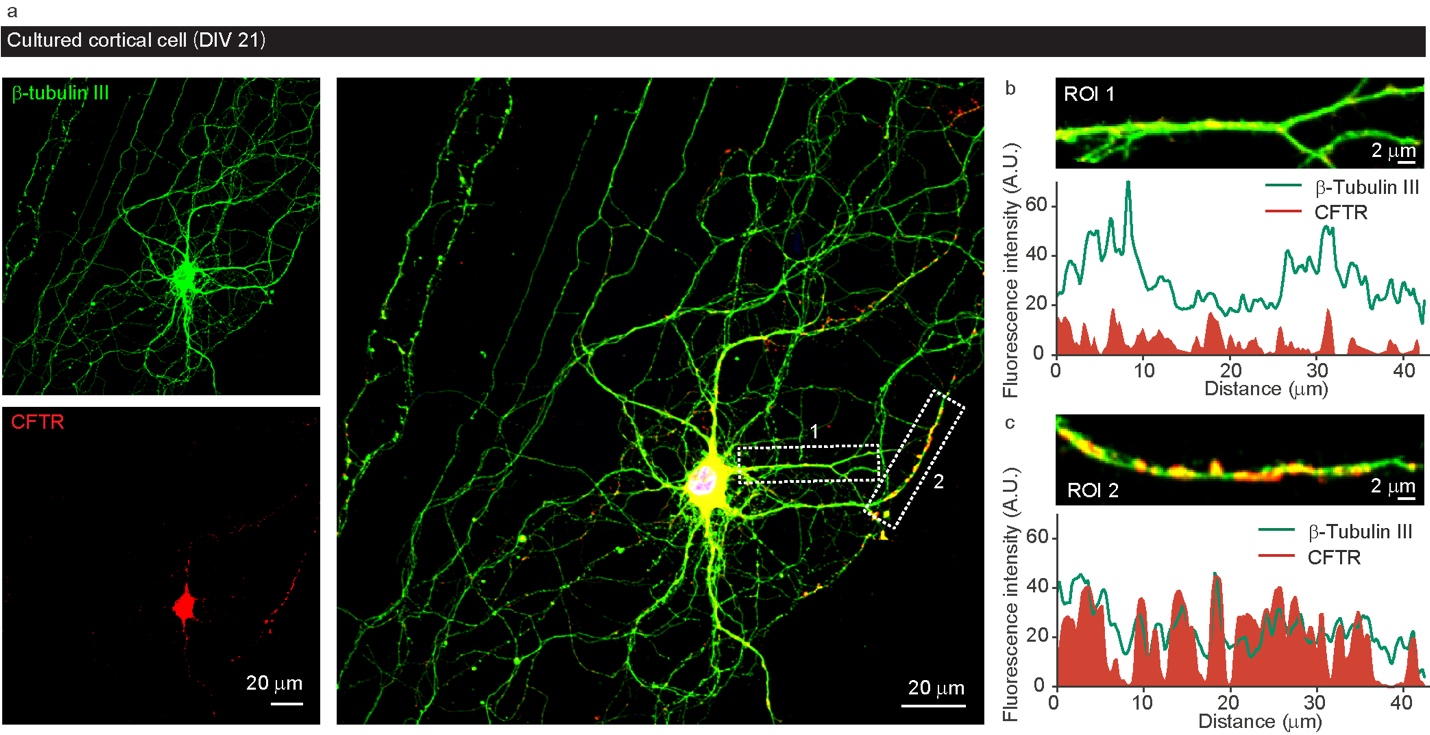

Supplementary Figure 4 | Expression of CFTR in mature cultured cortical neurons *in vitro*. Double immunolabeling of CFTR and neuron-specific tubulin III. CFTR was positively distributed in the cell body and some processes of neurons.

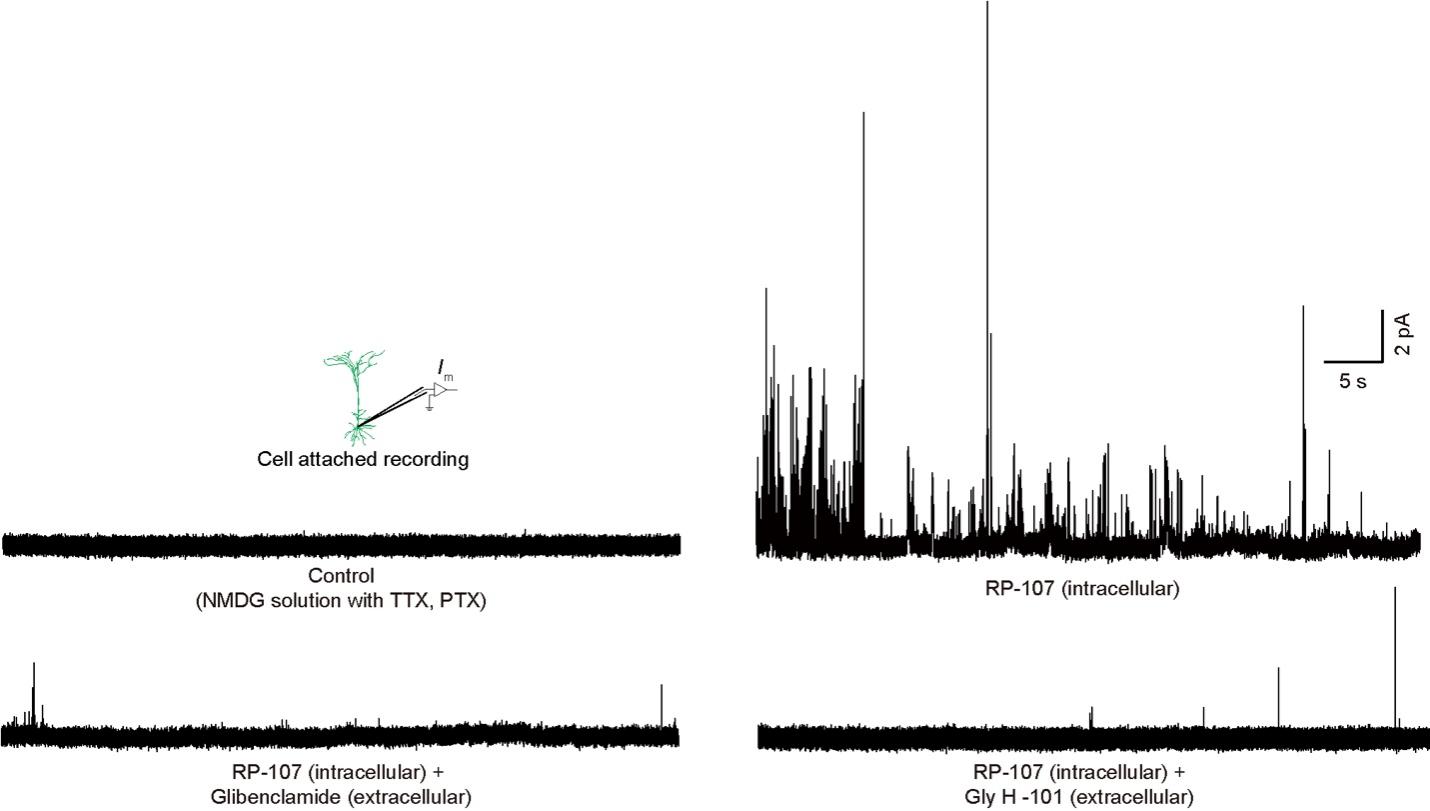

Supplementary Figure 5 | Activating CFTR increased the chloride current recorded by the cell-attached patch-clamp and could be eliminated by extracellular CFTR blockers.

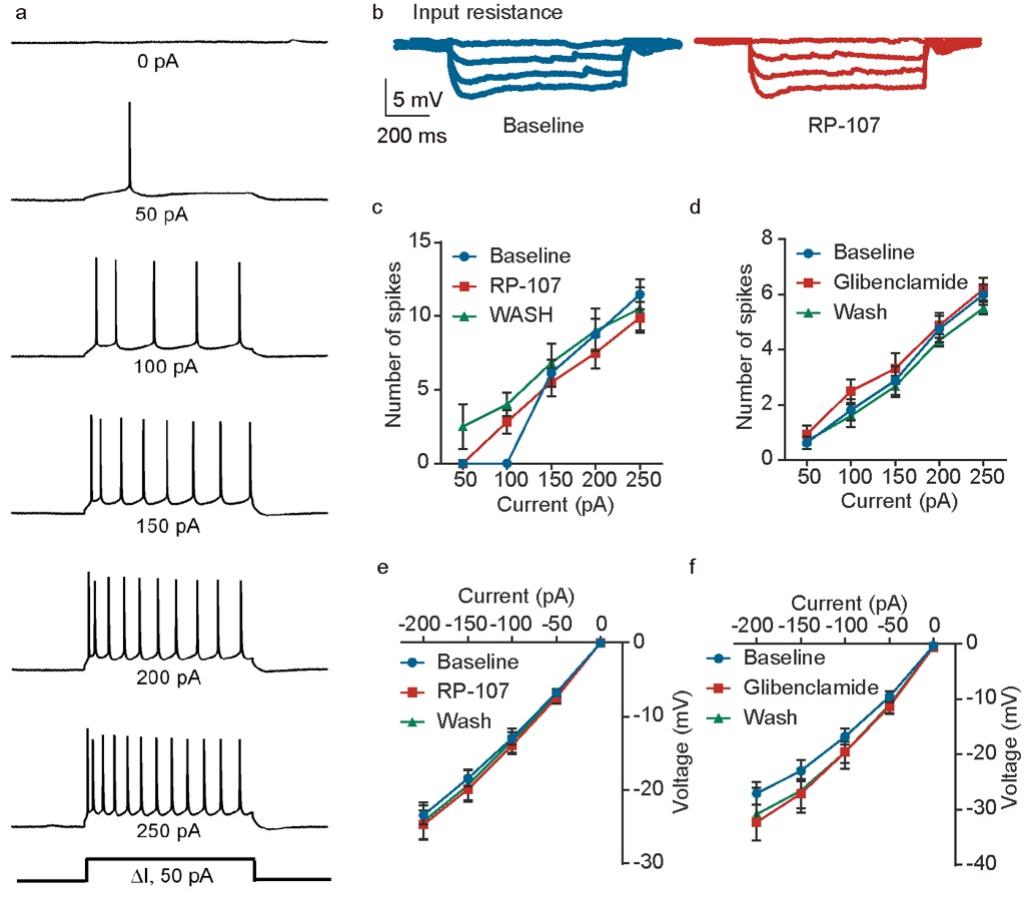

Supplementary Figure 6 | The CFTR channel has no significant effect on the number of evoked action potentials or input resistance. a. 50 pA step positive current could induce frequency-adaptive action potentials in layer V pyramidal neurons. b. Injected with different currents to measure the input resistance of neurons. c d. There is no significant difference in the number of neuronal evoked action potentials after CFTR activation or inhibition. e f. There is no significant difference in the input resistance of the neuron.

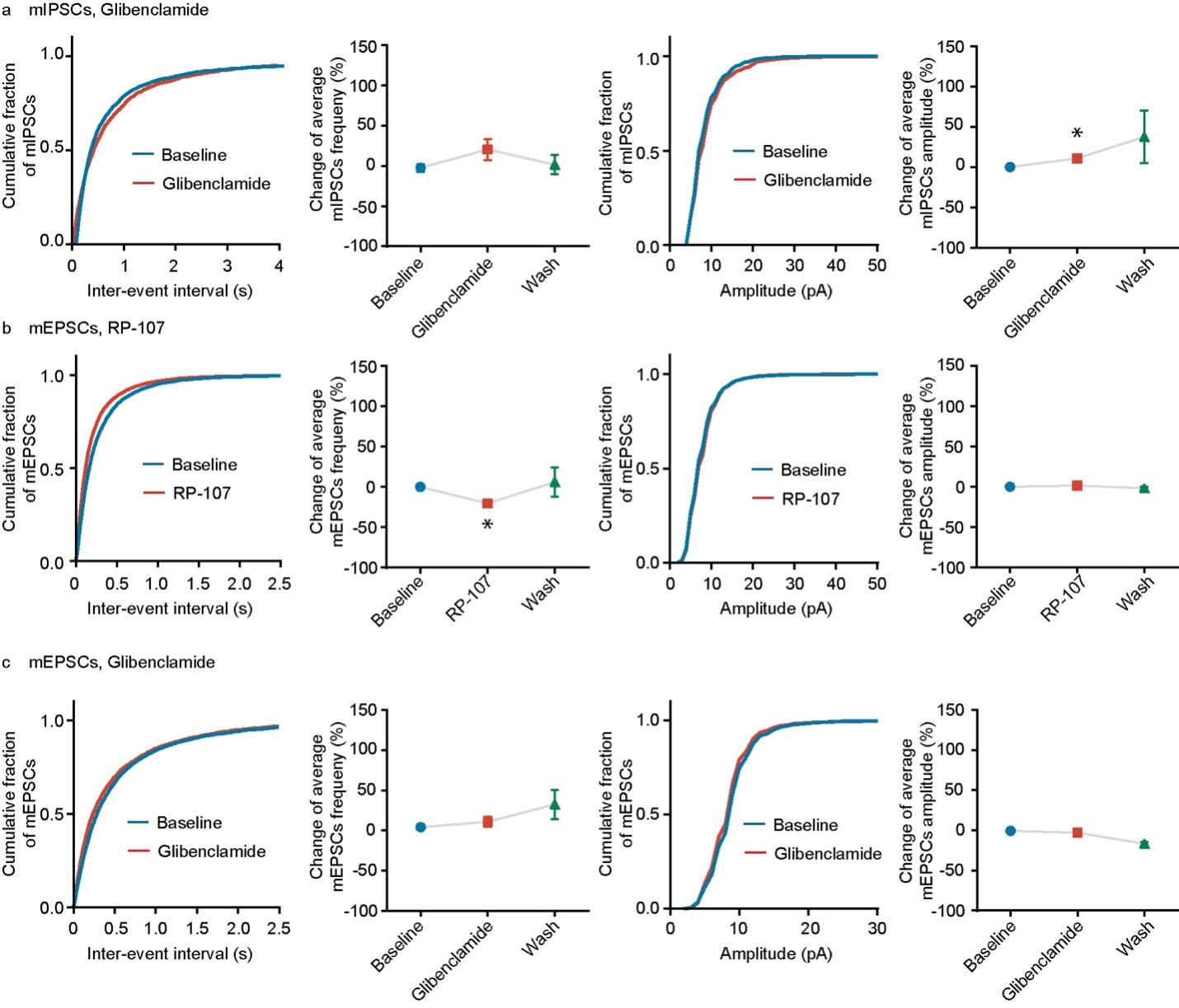

Supplementary Figure 7 | The effect of CFTR activity on miniature postsynaptic transmission. a, The effect of CFTR blocker Glibenclamide on mIPSC. b c, Activation (RP-107) or inhibition (Glibenclamide) of CFTR did not significantly change mEPSCs.

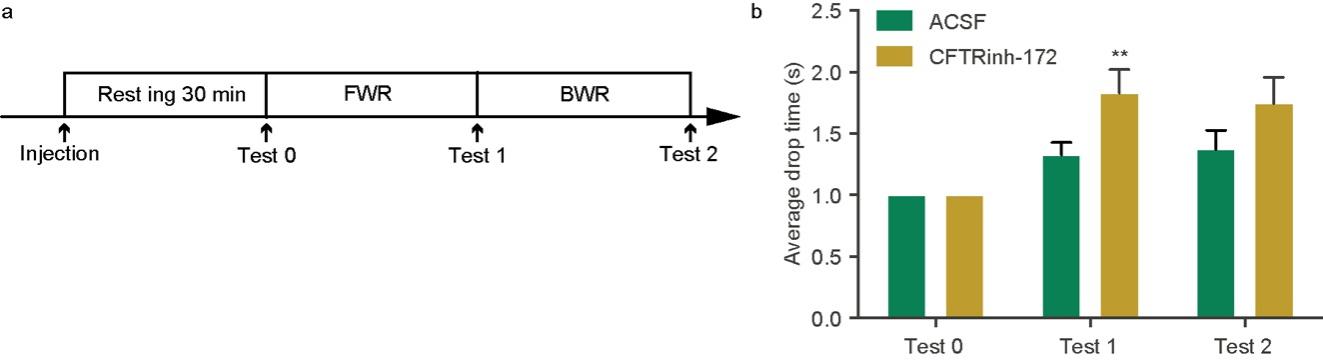

Supplementary Figure 8 | Inhibition of CFTR in the deep layer of the primary motor cortex enhanced the learning process of mice. a. After injecting the specific inhibitor CFTRinh-172 into the deep layer of the primary motor cortex on both sides, the mouse was trained to learn motor skills through rotarod training. b. The performance of mice after inhibiting CFTR was better than that of the ACSF group.

**Results of statistical test**
